# Hippocampal transcriptome analysis following maternal separation implicates altered RNA processing in a mouse model of fetal alcohol spectrum disorder

**DOI:** 10.1101/685586

**Authors:** Bonnie LJ Alberry, Christina A Castellani, Shiva M Singh

## Abstract

Fetal alcohol spectrum disorders (FASD) are common, seen in 1-5% of the population in the United States and Canada. Children diagnosed with FASD are not likely to remain with their biological parents, facing early maternal separation and foster placements throughout childhood. We have modeled FASD in mice via prenatal alcohol exposure and further induce early life stress through maternal separation. We report an association between adult hippocampal gene expression and prenatal ethanol exposure followed by postnatal separation stress that is related to behavioral changes. Clustering of expression profiles through weighted gene co-expression network analysis (WGCNA) identifies a set of transcripts, module 19, associated with anxiety-like behavior (*r* = 0.79, *p* = 0.002) as well as treatment group (*r* = 0.68, *p* = 0.015). Genes in this module are overrepresented by genes involved in transcriptional regulation and other pathways related to neurodevelopment. Interestingly, one member of this module, *Polr2a*, polymerase (RNA) II (DNA directed) polypeptide A, is downregulated by the combination of prenatal ethanol and postnatal stress in an RNA-Seq experiment and qPCR validation (q = 2e-12, p = 0.004, respectively). Together, transcriptional control in the hippocampus is implicated as a potential underlying mechanism leading to anxiety-like behavior via environmental insults. Further research is required to elucidate the mechanism involved and use this insight towards early diagnosis and amelioration strategies involving children born with FASD.

**SUMMARY STATEMENT:** Mouse hippocampal gene expression alterations following prenatal alcohol exposure and maternal separation are associated with behavioral deficits. Transcriptomic analysis implicates systems defect involving RNA processing, specifically including downregulation of *Polr2a*.

## INTRODUCTION

Ethanol is a teratogen that crosses the placenta and the blood-brain barrier, disrupting development. Alcohol use during gestation has been associated with various undesirable outcomes, including stillbirth (Cornman-Homonoff et al., 2012), spontaneous abortion (Kesmodel et al., 2002), premature birth (Sokol et al., 2007), and growth retardation (Sabra et al., 2018). Fetal alcohol spectrum disorder (FASD) is a direct result of gestational alcohol use, characterized by prenatal and postnatal growth restrictions, facial abnormalities, structural brain abnormalities, microcephaly, developmental delays, intellectual impairment, and behavioral difficulties (Chudley et al., 2005). Despite societal efforts to raise awareness of these risks, gestational alcohol use persists in North America. In Canada, an estimated 10% of pregnant women consume alcohol (Popova et al., 2017), and the prevalence of FASD in Canadian 7-9 year-olds is estimated between 2-3% (Popova et al., 2018). Similarly, in a cross-sectional study of four communities in the United States, the estimated prevalence of FASD ranged from 1.1-5% (May et al., 2018).The annual cost of FASD in Canada is estimated at approximately $1.8 billion, including costs due to productivity losses, the correctional system, and health care (Popova et al., 2016). While preventable, FASD remains a common and costly societal burden throughout an affected individual’s lifetime.

Children with FASD represent a significant proportion of children entering child care systems such as foster care or orphanages (Lange et al., 2013). Compared to a global estimate, the prevalence of FASD for children in care systems is 5.2-67.7 times higher in Canada (Lange et al., 2017). An unstable home environment results in a variety of postnatal stresses, often involving stress caused by maternal separation. In fact, exposure to early life stress via neglect or abuse increases the risk of psychiatric disorders later in life (Kisely et al., 2018). In rodents, early life stress also has a notable effect on hippocampus-specific learning and memory processes (Oomen et al., 2010; Pillai et al., 2018; Rice et al., 2008). Stress activates hippocampal neurons, ultimately leading to increased glucocorticoid receptor signaling (Hatalski et al., 2000). The hippocampus is critical for spatial learning and memory, through synaptic plasticity it is susceptible to the environment in ways that are often adaptive, although also make it vulnerable to chronic stress. As it stands, little is known about the combination of prenatal alcohol exposure and early life stresses experienced by children with FASD (Price et al., 2017). Children exposed to alcohol in utero and maltreatment during development are more likely to have impaired speech (Coggins et al., 2007), memory, attention, intelligence and other behavioral deficits (Henry et al., 2007; Koponen et al., 2009; Koponen et al., 2013). The molecular mechanisms involved in this interaction, however, have not been investigated.

Performing detailed molecular research on these topics in humans is challenging, as such we utilize animal models. Given that children with prenatal alcohol exposure often face postnatal chronic stress, we developed an animal model of FASD using C57BL/6J (B6) mice (Kleiber et al., 2011). Further, we have used this model to explore how postnatal stresses associated with maternal separation may compound behavioral and developmental deficits in mice following prenatal alcohol exposure (Alberry and Singh, 2016). The results follow the literature and show that pups prenatally exposed to ethanol develop learning deficits, anxiety-like behaviors, and changes in activity (Allan et al., 2003; Kaminen-Ahola et al., 2010; Kleiber et al., 2011; Marjonen et al., 2015). Specifically, in the Barnes Maze test for learning and memory, following prenatal alcohol exposure and postnatal separation stress, mice were slower to reach the location of a learned target (Alberry and Singh, 2016). In the open field test (OFT), mice are placed in a novel environment to freely explore and exploration of the center zone is indicative of anxiety-like behavior. In this test, mice prenatally exposed to alcohol are quicker to enter the center zone than controls. Finally, the home cage activity test (HC) is used to assess activity in a familiar environment. In the HC test, mice that had faced postnatal maternal separation were less active than controls (Alberry and Singh, 2016). Also, early life stress introduced by maternal separation and isolation during early development in mice may lead to increased anxiety-like behaviors in adults (Romeo et al., 2003). The results of rodent models of maternal separation have found the first 10 postnatal days represent the most critical period (Fenoglio et al., 2006), as such, separation models have focused on this time (Franklin et al., 2010; Veenema et al., 2008).

Neurodevelopment lasts into adulthood and can be affected by ethanol exposures and external stresses at anytime during this period, suggesting that early postnatal environment can alter adult behaviour in progeny that had faced prenatal ethanol exposure (Chokroborty-Hoque et al., 2014). This influence may involve changes in gene expression as the foundation for alterations in neurodevelopment and brain function. Here, we use our mouse model to identify changes in hippocampal gene expression following prenatal ethanol exposure and postnatal maternal separation stress in mice.

## RESULTS

RNA-Seq was performed on hippocampal RNA samples from three individuals for each of four groups of mice representing a control group with no experimental interventions, an ethanol group of mice prenatally exposed to ethanol, a stress group with mice subjected to postnatal maternal separation stress, and an ethanol + stress group with mice prenatally exposed to ethanol followed by postnatal maternal separation stress. Transcriptomes from these 12 samples were assessed via RNA-Seq to determine how they differ between treatment groups and control, with sample hierarchical clustering indicating no obvious outliers (Supplementary Figure 1). Weighted gene co-expression network analysis (WGCNA) was used to cluster transcripts into modules based on correlated expression across samples that can be assessed in relation to other known traits. For optimal correlations between modules and traits of interest, we chose a cutHeight of 0.35. A lower cutHeight results in more modules with fewer genes per module, with adjacent modules sharing trait associations, while increasing cutHeight results in fewer, larger modules. Here, it results in 44 modules, represented by 43 to 11 946 transcripts in each module (Figure 1A).

**Figure 1.**
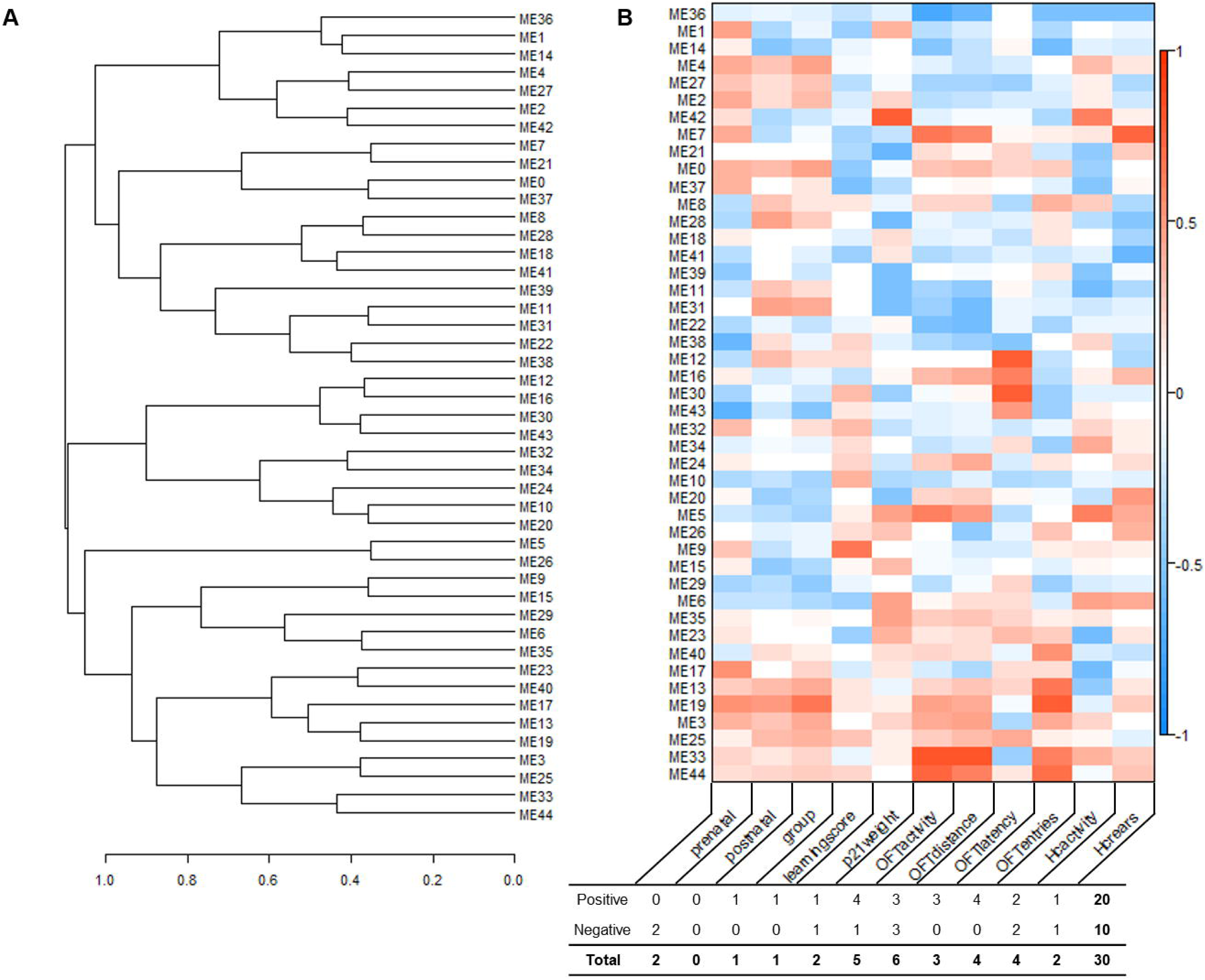
Module formation and trait association using weighted gene co-expression network analysis (WGCNA). (A) Hierarchical clustering dendrogram of module eigengenes with dissimilarity of eigengenes given by 1-(eigengene correlation). (B) Module-trait correlation heatmap for 11 traits: prenatal or postnatal treatment, experimental group (group), Barnes maze learning score (learningscore), weight at postnatal day 21 (p21weight), open field test activity (OFTactivity), distance (OFTdistance), latency to enter the center (OFTlatency), number of center entries (OFTentries), home cage activity (HCactivity), and number of rears (HCrears), with positive (red) and negative (blue) correlations, with the number of significantly (p < 0.05) positively or negatively correlated modules with each trait indicated below.

### WGCNA reveals associations between gene modules and treatment outcomes

We assessed the correlation between each module produced using WGCNA and 11 traits specified either by assignment or previous measurement, including prenatal treatment, postnatal treatment, treatment group, Barnes Maze learning score (learningscore), weight on postnatal day 21 (p21weight), open field test measures of activity (OFTactivity), distance (OFTdistance), latency to enter the center (OFTlatency), and number of entries to the center (OFTentries), as well as home cage measures of activity (HCactivity) and number of rears (HCrears) (Alberry and Singh, 2016) (Supplementary Table 1). These traits have unique module correlations with visibly shared patterns (Figure 1B). Although some modules share similar correlations, no two traits have the same correlation status with each module. Modules clustering next to one another share similar patterns of correlation between some traits, but no two modules have the same profile at this level. Using a nominally significant cutoff (p < 0.05), 30 module-trait relationships emerge, with a range from zero to six significantly correlated modules per trait. Further, most of these relationships (20/30) are positive correlations, while others (10/30) are negative. Also, 20 of the 44 modules (45.56%) are significantly correlated to at least one trait, four correlated to two traits, and three modules correlated to three traits. It is worth noting, however, that these traits are not independent. While there are several overlapping correlations between traits, only one module, module 19 (ME19) is correlated with both experimental treatment and measured behavioral outcome.

### Module 19: RNA polymerase II-associated functions are correlated to experimental group and anxiety-like behavior

One module, ME19, is correlated with experimental group (group) (r = 0.68, p = 0.015), as well as number of entries into the center during the open field test (OFTentries) (r = 0.79, p = 0.002). This module is composed of 895 transcripts that align to 739 annotated genes, including two complete protein complexes (Supplementary Table 2). Epidermal growth factor and its receptor (EGF:EGFR) are represented in this module by *Egf* and *Egfr* transcripts. In addition, the N-methyl-D-aspartate (NMDA) glutamate receptor (NMDAR) is represented by *Grin2b*, glutamate receptor, ionotropic, NMDA2B (epsilon 2), and *Grin1*, glutamate receptor, ionotropic, NMDA1 (zeta 1). Using a modest threshold for inclusion (p < 0.05), Module 19 genes implicate KEGG pathways important in transcription such as RNA degradation (p = 0.006), as well as neurodevelopment and neurodegeneration, including adherens junction (p = 0.015) (Table 1, Supplementary Table 2). Major gene ontologies implicated by genes in this module using the same threshold are important for neurodevelopment and neurodegeneration, such as beta-catenin-TCF complex (p = 6 × 10^−4^), Notch signaling (p = 7 × 10^−4^), and MAPK cascade (p = 9 × 10^−4^), as well as transcription, including RNA polymerase II functions (p = 0.001) (Table 1, Supplementary Table 2).

**Table 1.** Top 5 most significantly over-represented KEGG pathways and gene ontology (GO) terms represented by genes in module 19.

| Term | p-value |
| --- | --- |
| **KEGG pathway** |  |
| RNA degradation | 0.0051 |
| Rap1 signaling | 0.0079 |
| Prostate cancer | 0.0088 |
| Arginine and proline metabolism | 0.0102 |
| Adherens junction | 0.0111 |
| **Cellular components** |  |
| beta-catenin-TCF complex | 0.0006 |
| Melanosome | 0.0009 |
| pigment granule | 0.0009 |
| azurophil granule lumen | 0.0047 |
| Microbody | 0.0055 |
| **Biological processes** |  |
| positive regulation of binding | 0.0006 |
| Notch signaling pathway | 0.0007 |
| MAPK cascade | 0.0009 |
| regulation of protein targeting to mitochondrion | 0.0011 |
| regulation of protein metabolic process | 0.0012 |
| **Molecular functions** |  |
| RNA polymerase II regulatory region sequence-specific DNA binding | 0.0010 |
| chromo shadow domain binding | 0.0014 |
| RNA polymerase II distal enhancer sequence-specific DNA binding | 0.0025 |
| ubiquitin-like protein ligase binding | 0.0046 |
| RNA polymerase II core promoter proximal region sequence-specific DNA binding | 0.0057 |

### Prenatal ethanol exposure and early life maternal separation stress are associated with changes in gene expression

Differentially expressed genes were determined at the transcript level for each experimental treatment group compared to the control group (Supplementary Table 3). Genes with larger effect size (beta values) tend to also reach greater significance (Figure 2A, B, C). Filtering for existing transcripts aligned to the mouse genome (mm10), 164 unique transcripts were implicated by ethanol, 116 by stress, and 217 by the combination of two treatments (*p* < 0.01) (Figure 2D). There was some overlap between lists, with 13 transcripts shared by all three lists. Differentially expressed genes for each treatment group were analyzed for enrichment of gene ontology and KEGG pathways using annotated genes from these transcripts (Table 2, Supplementary Table 3). Transcripts differentially expressed following prenatal ethanol treatment are important for mRNA processing (*p* < 1.01 × 10^−5^) and synapse localization (*p* < 0.001). Following postnatal maternal separation stress, transcripts important for cell polarity (*p* = 8.64 × 10^−5^) and several neurological pathways (*p* < 0.001) are differentially expressed. For mice exposed to both prenatal ethanol and postnatal stress, altered transcripts are important for stimulus response (*p* < 2.39 × 10^−4^). Taken together, some ontologies and pathways altered by either prenatal ethanol exposure or postnatal stress are shared, particularly with synaptic functions (synapse, synapse part, postsynapse, GABAergic synapse). In addition, genes related to hemoglobin binding have altered expression in the Ethanol as well as Ethanol + Stress groups (p < 1.69 × 10^−5^).

**Table 2.**
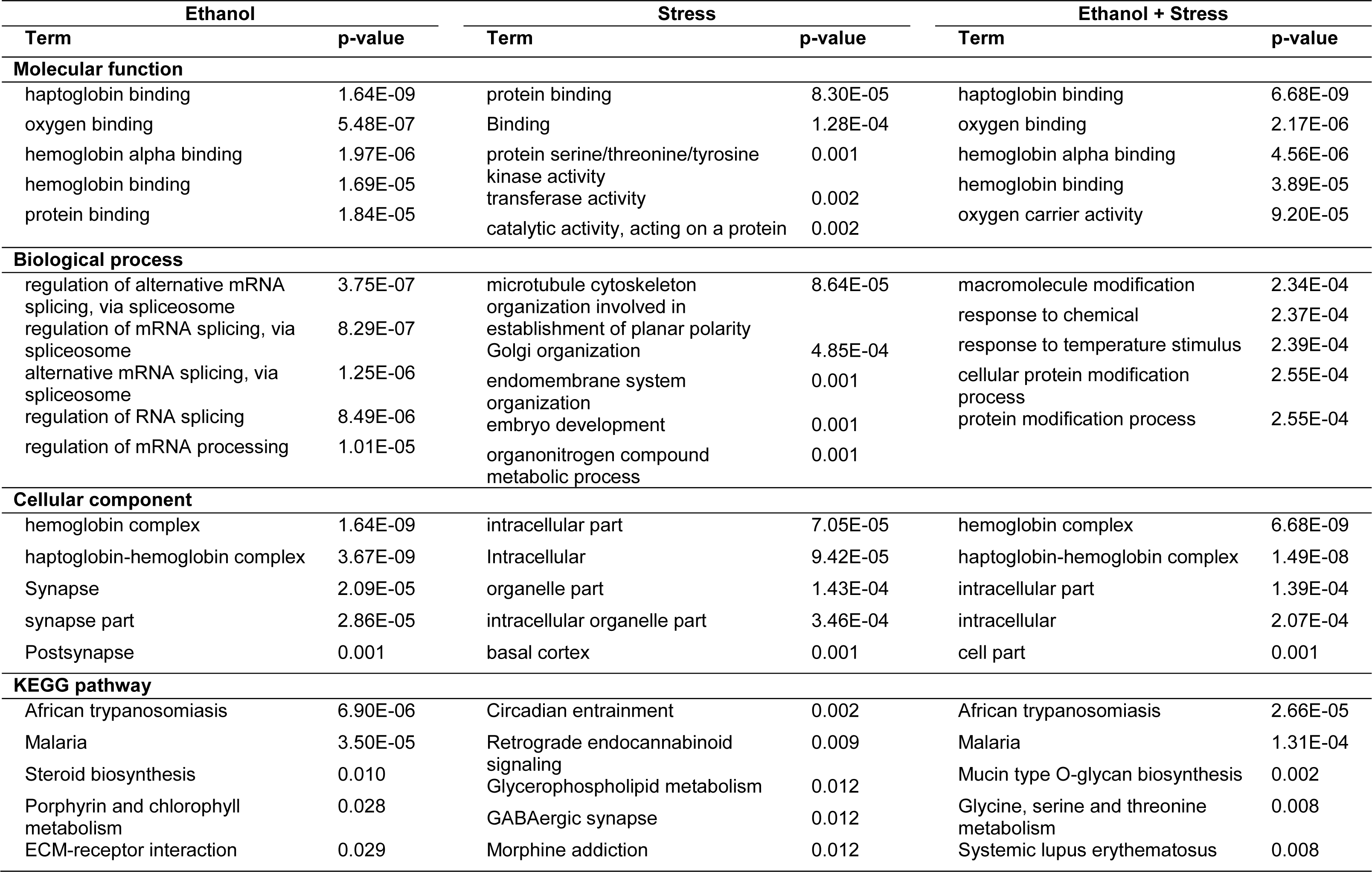
Top 5 most significantly over-represented gene ontology (GO) terms and KEGG pathways represented by annotated genes of transcripts differentially expressed in the Ethanol, Stress, or Ethanol + Stress groups compared to control (p < 0.01).

**Figure 2.**
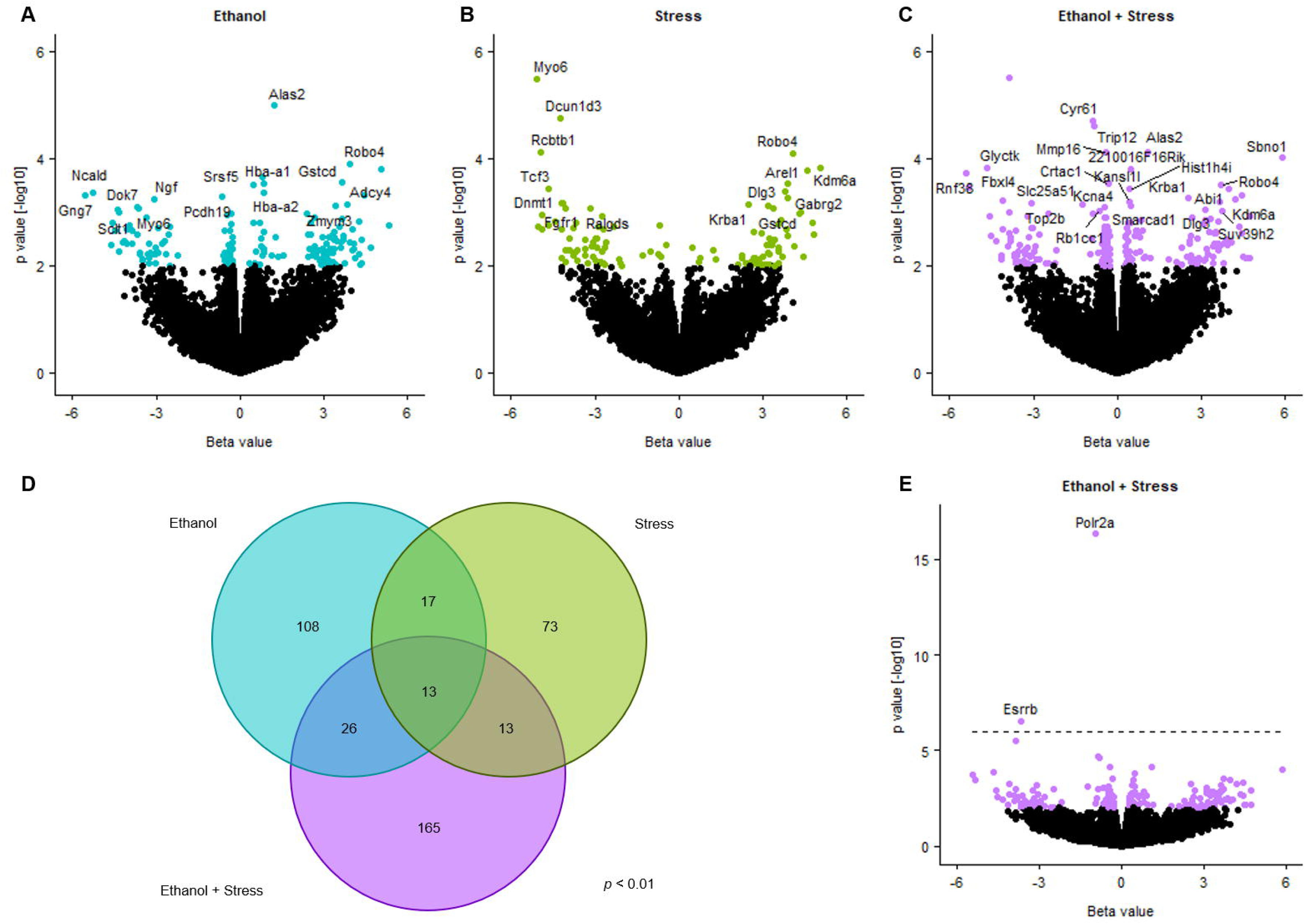
Differential gene expression between groups as detected by sleuth. Volcano plots indicating effect size (Beta value) and significance (*p* < 0.01 by color) for each transcript in Ethanol (A), Stress (B), and Ethanol + Stress (C) groups compared to control treatment with nominally significant (*p* < 0.001), annotated genes labeled. Venn diagram of overlapping transcripts differentially expressed in each treatment as compared to controls (*p* < 0.01, D). (E) Volcano plot indicating effect size (Beta value) and significance for transcripts with an expanded scale and significant (*q* < 0.05) annotated genes labelled for Ethanol + Stress compared to control.

Some overlap exists between these three lists, specifically when examining the top 25 most significant annotated transcripts in each list (Table 3). Most notably, *Robo4*, Roundabout guidance receptor 4, and *Krba1*, KRAB-A domain containing 1, are upregulated in each treatment group, with effect sizes ranging from 2.40 to 4.07. *Alas2*, aminolevulinic acid synthase 2, erythroid, is the most significantly differentially expressed gene following ethanol treatment (beta = 1.21, p = 1.04 × 10^−5^), and is also upregulated in the Ethanol + Stress group (beta = 1.08, p = 7.71 × 10^−5^). Similarly, *Suv39h2*, suppressor of variegation 3-9 2, is also upregulated following prenatal ethanol exposure (beta = 3.64, p = 1.34 × 10^−3^) as well as following the combined Ethanol + Stress treatments (beta = 3.74, p = 9.76 × 10^−4^). *Kdm6a*, lysine (K)-specific demethylase 6a, *Sbno1*, strawberry notch 1, and *Dlg3*, discs large MAGUK scaffold protein 3, are shared between the Stress and Ethanol + Stress groups as upregulated when compared to controls.

**Table 3.**
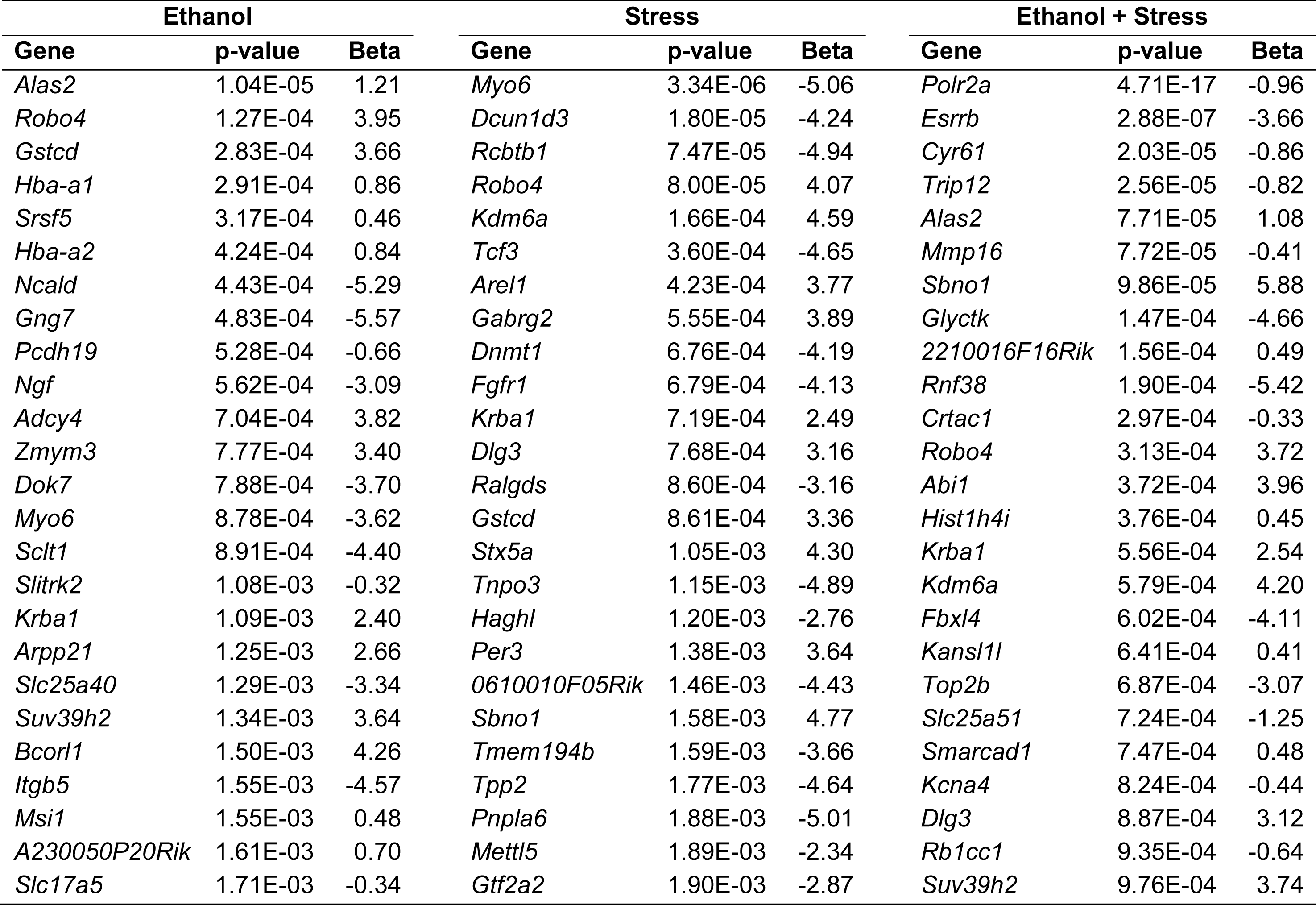
Top 25 annotated gene transcripts identified in each treatment group, where beta value represents effect size for each transcript detected.

Two differentially expressed genes show robust changes in differential expression between the ethanol + stress group compared to the control group, adjusted for a stringent false discovery rate (q < 0.05), *Esrrb*, estrogen related receptor, beta and *Polr2a*, polymerase (RNA) II (DNA directed) polypeptide A (Figure 2D). Each of these is the result of a single transcript downregulated following the combination of Ethanol + Stress (Figure 3A, B). Interestingly, these two genes are also members of Module 19 from WGCNA. The low level of *Esrrb* expression in the RNA-Seq experiment excluded it from further validation. Downregulation of *Polr2a* was validated by qPCR displaying 1.28-fold decrease in expression following Stress (*p* = 0.048), and 1.59-fold decrease following the combination of Ethanol + Stress (*p* = 0.004) (Figure 3C).

**Figure 3.**
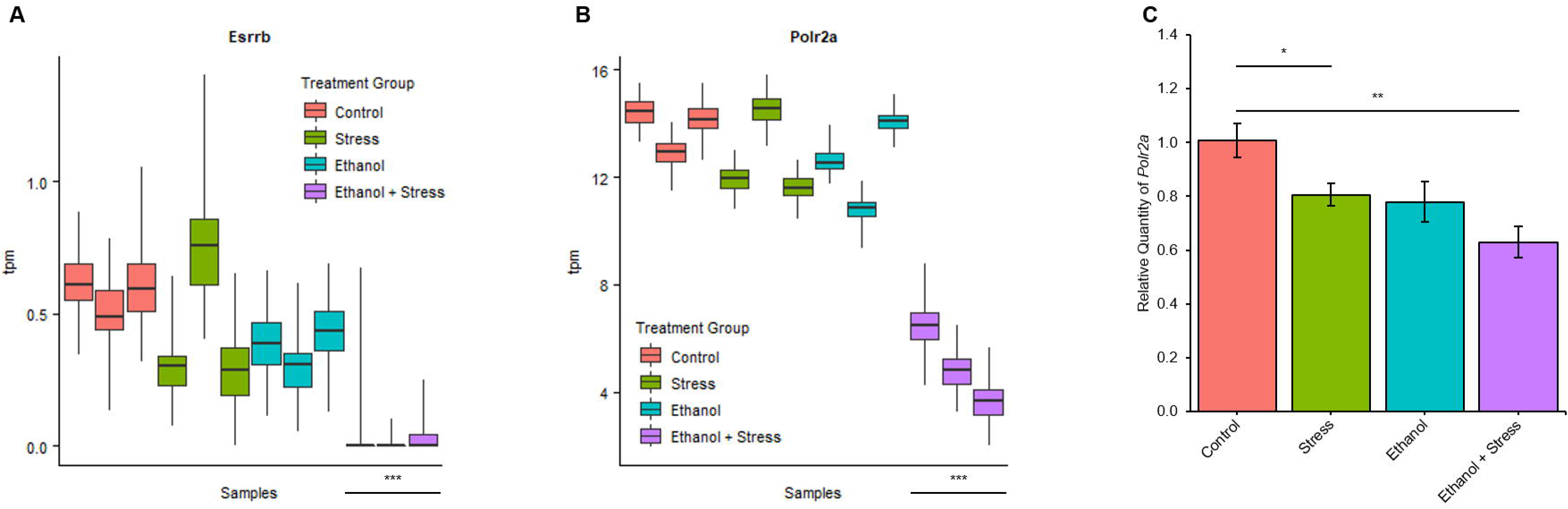
Transcript-specific differential gene expression following Ethanol + Stress treatment. *Esrrb* (A) and *Polr2a* (B) transcript abundance in transcripts per million (tpm ± inferential variance) as detected by sleuth from RNA-Seq, ***q-value < 0.05. (C) Relative quantity (± s.e.m.) of *Polr2a* is decreased 1.24 -fold with postnatal stress alone, and 1.59-fold when mice were prenatally exposed to ethanol and postnatal stress, as detected by reverse transcription qPCR, *p < 0.05, **p < 0.01.

## DISCUSSION

FASD is a complex societal burden that while preventable, remains common. Besides prenatal exposure to alcohol, children born with FASD are often also exposed to stressful postnatal upbringing that invariably includes early maternal separation. The main objective with this study was to understand how early life stresses may complicate molecular changes and behavioral deficits following exposure to prenatal alcohol. The experimental design used has previously assessed the behavioral changes that result from prenatal alcohol exposure and early postnatal stress(Alberry and Singh, 2016), while the molecular changes reported here are focused on adult hippocampal transcriptome. The results show that the manifestation of FASD-related deficits may be compounded by additional maternal separation stress via alterations in hippocampal gene expression. Specifically, the analysis of this multifaceted data by WGCNA has led to the identification of modules of genes similarly expressed between samples that correlate with treatment and behavioral outcomes. It has identified one module of genes, module 19, that correlates with treatment and an anxiety-like behavior in the progeny (number of center entries in the open field test). Genes in this module are related to transcription and neurodevelopment through various gene ontologies and KEGG pathways. We also performed a transcript level differential expression analysis, finding that early life stress as well as the combination of prenatal ethanol exposure and postnatal stress result in downregulation of a transcript responsible for an RNA polymerase II subunit (*Polr2a*) in the hippocampus. Together, these results suggest that changes in behavior following prenatal alcohol exposure and postnatal stress likely result from altered hippocampal gene expression.

### WGCNA reveals FASD-relevant module of co-expressed transcripts

We have previously described behavioral changes in these mice as significantly different between treatment groups (Alberry and Singh, 2016), and here add to that data through the analysis of RNA-Seq gene expression results from the adult hippocampus of the same mice as well as the novel use of WGCNA in an *in vivo* FASD model. WGCNA is a valuable systems biology tool for determination of modules of correlated gene expression and relating these modules to sample traits (Langfelder and Horvath, 2008). WGCNA had identified a single module of hippocampal gene expression, module 19, that is correlated with experimental treatment as well as with a reliably measured phenotypic outcome. In this case, the number of entries into the center zone of the open field test by mice is analogous to an anxiety-like behavior (Choleris et al., 2001; Prut and Belzung, 2003). Interestingly, several FASD-related deficits, including learning, memory, stress response and anxiety-like behaviors are under hippocampal control. Specifically, increased adult hippocampal neurogenesis has been associated with reduced anxiety-like behavior in mice (Hill et al., 2015), and stress-induced alterations in hippocampal microglia have also been associated with anxiety-like behavior (Kreisel et al., 2014). In addition, exposure to chronic stress is associated with activated hippocampal microglia in rats (Tynan et al., 2010). Such results argue that the WGCNA analysis has identified a set of genes represented in module 19 that are relevant to FASD etiology.

Next, we focused on the set of genes represented in module 19. We argue that the genes affected represent pathways that may be disturbed in FASD. Genes responsible for the MAPK cascade (21 genes) and notch signaling (8 genes) are included in this module. MAPK signaling has been identified as a strong candidate pathway in FASD and is fundamental to fetal development (Lombard et al., 2007). MAPK signaling cascade has also been implicated in ethanol-induced apoptosis through activation of p53 signaling in neural crest cells (Yuan et al., 2017). In addition, human neural precursors display impaired neurogenesis following alcohol exposure via downregulation of MAPK genes (Louis et al., 2018). The MAPK pathway is also involved in chronic stress, with increased negative regulation via MAPK phosphatase-1 in the hippocampus of chronically stressed rodents (Duric et al., 2010). Together, it is unsurprising to find the MAPK cascade implicated by gene represented in Module 19, with expression correlated to prenatal alcohol exposure and maternal separation stress as well as anxiety-like behavior in the resulting progeny. Further, genes involved in Notch signaling are also overrepresented in module 19. Notch signaling is important for neurogenesis and embryonic development through its influence on expression of associated transcription factors, even in the adult brain (Ables et al., 2011). Notch signaling has also been reported as altered in a number of prenatal alcohol exposure models involving mice (Ninh et al., 2019) and zebrafish (Muralidharan et al., 2018). Our results further support the notion that MAPK and notch signaling, alongside other signaling cascades, implicate a role for altered transcriptional control in the adult hippocampus following prenatal alcohol exposure and early life stress.

### Gene expression patterns implicate neurodevelopmental dysregulation in FASD

To the best of our knowledge, this is the first report of genome-wide changes in hippocampal gene expression following a continuous preference exposure of alcohol in a mouse model of FASD. Our results argue that prenatal ethanol exposure and early maternal separation lead to alterations in specific hippocampal genes depending on the type and combination of exposures. Of interest to this discussion are two transcripts, *Robo4* and *Krba1*, that are among the top 25 most significantly altered transcripts in each treatment group, both upregulated in every group compared to control (p < 0.001). The protein product for *Robo4* is the roundabout (Robo) receptor, with interacts with the astrocyte-secreted slit guidance ligand 2 (Slit2) during central nervous system development, specifically as an axon repellant during axon guidance (Brose et al., 1999). *Krba1* may be relevant as a transcription factor, it is predicted to regulate transcription via DNA template binding as a zinc-finger protein that represses RNA polymerase promoters (Urrutia, 2003). Gestational long-term exposure to air pollution (measured as particulate matter with a diameter < 2.5 µm) was associated with increased cord blood *KRBA1* expression in females (Winckelmans et al., 2017). If upregulation of these two genes following each of our treatments results in decreased viability, abnormal synapse formation, and improper regulation of transcription during neurodevelopment, they may have the potential to contribute to observed behavioral deficits in FASD.

We also note that two transcripts, *Alas2* and *Suv39h2*, are among the top 25 most significantly altered transcripts in two treatment groups (Ethanol and Ethanol + Stress). There have been mixed reports regarding alcohol exposure and *Alas2* expression. In rat placenta following voluntary maternal ethanol consumption, *Alas2* is downregulated (Rosenberg et al., 2010). In a postnatal lead exposure model, *Alas2* is upregulated in the hippocampus following lead exposure in male rats (Schneider et al., 2012). *Alas2* is also downregulated in whole blood one hour after a stress exposure in heavy drinkers, characterized by regular alcohol use (Beech et al., 2014). In a mouse model of chronic social stress, Alas2 was upregulated in the prefrontal cortex after 13 days of repeated stress (Stankiewicz et al., 2014). The protein product of *Suv39h2* is a histone methyltransferase, specifically for H3K9, often resulting in trimethylation leading to inhibition of gene expression (Peters et al., 2001). Differential histone methylation in the brain has also been implicated in FASD (Chater-Diehl et al., 2016; Chater-Diehl et al., 2017).

Finally, three transcripts, *Kdm6a, Sbno1*, and *Dlg3*, were among the top 25 most significantly altered transcripts in response to the stress treatments (Stress and Ethanol + Stress groups). *Kdm6a* encodes a histone demethylase that removes suppressive chromatin marks. When forebrain microglia were exposed to dying cells in vitro, expression of this histone demethylase increased, with research suggesting it controls gene expression in microglia for clearance of dying neurons and non-functional synapses (Ayata et al., 2018). *Sbno1* is a nuclear localized transcriptional regulator important in Notch and Hippo signaling (Watanabe et al., 2017). In a WGCNA analysis of gene expression from post-mortem prefrontal cortex of patients with schizophrenia, *Sbno1* was in the lone module associated with prior schizophrenia GWAS loci (Fromer et al., 2016), results that were independently replicated (Radulescu et al., 2018), suggesting its expression patterns may relate to brain function. *Dlg3* encodes synapse-associated protein 102 and is important for synapse formation in the brain. *Dlg3* knockout mice have synaptic plasticity impairments, including impaired spatial learning (Cuthbert et al., 2007), and mutations in DLG3 cause non-syndromic X-linked intellectual disability (Tarpey et al., 2004). We report upregulation of a *Dlg3* transcript in the Stress and Ethanol + Stress groups compared to control. Other research has shown that DLG3 overexpression in a cancer cell culture model inhibits proliferation and induces apoptosis (Liu et al., 2014). Taken together, the upregulation of these three genes in the Stress and Ethanol + Stress groups may lead to altered brain transcriptional regulation and synaptic plasticity required for normal learning processes and brain function.

### Gene expression dysregulation compliments WGCNA results

The molecular functions most significantly enriched by Module 19 genes involve RNA polymerase II DNA binding. These are genes correlated with experimental treatment group and anxiety-like behavior. When looking for differentially expressed genes, we also find that the most significantly differentially expressed transcript coding for an RNA polymerase II subunit (*Polr2a*) is downregulated following stress and ethanol + stress as compared to control. *Polr2a*, polymerase (RNA) II (DNA directed) polypeptide A, encodes for the largest subunit of RNA polymerase II, an enzyme responsible for mRNA synthesis. Like the combination of ethanol + stress we present here, a mouse model of stress + morphine from postnatal days 5 to 9 found decreased *Polr2a* expression following stress alone (Juul et al., 2011). Conversely, when combined with morphine treatment, there was an associated upregulation of *Polr2a*, suggesting expression of this gene is particularly sensitive to environmental insults. Ours is not the first evidence of Polr2a alterations in FASD research, in a mouse model of binge-like prenatal ethanol exposure, increased promoter DNA methylation of *Polr2a* was found in the hippocampus (Chater-Diehl et al., 2016). Additionally, *Polr2a* expression is decreased in the dentate gyrus of the hippocampus alongside increased anxiety-like behavior following induced glucocorticoid receptor overexpression during the first 3 weeks of life as a model of early life adversity (Wei et al., 2012). WGCNA has also been used to create gene co-expression modules related to ethanol exposure in human embryonic stem cells to help understand molecular mechanisms underlying FASD. Despite vastly different experimental design, one module associated with ethanol treatment in the stem cells was also most significantly enriched with RNA polymerase II activity (Khalid et al., 2014).

In conclusion, we have examined RNA isolated from adult hippocampus, a more focused tissue type than whole brain homogenate, representing RNA from a mixed cell population. Multiple lines of evidence suggest microglia and neurons respond differently and need to be assessed separately. In our WGCNA analysis, Module 19 was associated with changes in hippocampal gene expression and anxiety-like behavior, an association potentially driven by microglia. Additionally, the *Kdm6a* histone demethylase found differentially expressed in stress and ethanol + stress groups is important for microglia function in clearance of dying neurons. In fact, developmental alcohol exposure leads to lasting disruption of microglia in rodents (Chastain et al., 2019). Further work in this area should focus on cell type differences that underlie the observations described here. Also, alterations in gene expression in this study were assessed at postnatal day 70, following prenatal ethanol exposure and/or postnatal stress up to postnatal day 14. It seems unlikely that the direct effect of the treatments used are persisting 8 weeks later. It is logical to argue that these exposures are causing long-lasting transcriptomic effects. In this context, epigenetic mechanisms are known to be involved in the programming of gene expression throughout development, and as mediators of experience to finely control expression. As in our results, alterations to epigenetic marks such as DNA methylation, histone modifications, and microRNA expression have been implicated following prenatal alcohol exposure and early life stress. As such, future experiments should aim to classify alterations in epigenetic terms that may be responsible for the persistent changes in gene expression seen over a long period of time. Finally, we have demonstrated that transcriptome changes persist in the hippocampus of adult mice prenatally exposed to alcohol and/or postnatally exposed to early life stress. Some transcripts are co-expressed in a way that correlates with experimental treatment and behavioral outcomes. These transcripts code for genes important in RNA processing and management throughout neurodevelopment and beyond. In addition, some transcripts are significantly differently expressed between treatment groups. Specifically, *Polr2a* is downregulated following early life stress as well as the combination of prenatal ethanol exposure and postnatal stress. These lasting alterations in transcripts responsible for RNA processing likely underlie the behavioral deficits observed in these mice. They suggest that postnatal stresses further complicate the effects caused by prenatal alcohol exposure in FASD. Further understanding into how these changes occur and persist may result in earlier detection, prognosis, and amelioration in humans faced with FASD.

## METHODS

### Animals

C57BL/6J (B6) mice (*mus musculus*) were obtained from Jackson Laboratories (Bar Harbor, ME, USA) and bred in the Animal Care Facility at Western University (London, Ontario, Canada). Mice were housed in same-sex colonies of up to four individuals with unrestricted access to food and water. Cage, bedding, and nest material were consistent between cages. The colony room operated on a 14:10 h light-dark cycle, with a humidity range from 40%-60%, and temperature range from 21-24 °C. Protocols complied with ethical standards established by the Canadian Council on Animal Care and approved by the Animal Use Subcommittee at Western University (London, Ontario, Canada).

### Continuous preference drinking model

Following the continuous preference drinking (CPD) model, 10-week old females were individually housed and randomly assigned to either the control group, dams with free access to water only, or the ethanol group, voluntary ethanol consumption dams with free access to water and a 10% ethanol in water solution (Alberry and Singh, 2016; Kleiber et al., 2011). The ethanol group were presented ethanol solution in water with increasing concentrations available from 2%, 5%, and 10%, each introduced after 48 hours of exposure to the previous concentration. Following 10% ethanol availability over a 10-day period, females were mated with 15-week-old males, with only water available. Males were removed after 24 h, representing gestational day 0. Ethanol was available to ethanol females until postnatal day 10. Ethanol was then removed and only water was provided to all females for the duration of the study. While blood alcohol levels were not taken to minimize maternal stress, voluntary consumption of 10% ethanol at 14 g ethanol per kg body weight per day has been shown to produce peak blood alcohol levels of 120 mg dl^−1^ (Allan et al., 2003). Experimental mice consumed an average of 8 g ethanol per kg body weight per day, representing a modest level of ethanol exposure. This level of exposure has produced modest behavioral deficits in these offspring, including hyperactivity in a novel environment, hypoactivity in a home cage environment, and learning deficits (Alberry and Singh, 2016).

### Early life stress via maternal separation & isolation

Early life stress via postnatal maternal separation occurred as previously described (Alberry and Singh, 2016; Benner et al., 2014; Savignac et al., 2011). Half of pups in each litter (2-4 pups per litter) were randomly selected for separation stress on postnatal day 2. Tail coloring with permanent marker was used to distinguish pups. Pups were removed and isolated for 3 h per day during the light phase from 10:00 – 13:00 up to and including postnatal day 14 in 8 oz. Dixie cups with bedding and nest material. Pups not selected for separation remained with the dam and other littermates during this time. Weaning occurred at postnatal day 21, with same-sex littermates housed in cages of 2-4 individuals. Following treatments, experimental mice belonged to one of four groups: control, ethanol, stress, or ethanol + stress.

### Hippocampal dissection & RNA isolation

On postnatal day 70, male mice were sacrificed via carbon dioxide asphyxiation and cervical dislocation. Hippocampus was dissected from whole brain as previously described (Spijker, 2011). Hippocampus samples were transferred to individual tubes containing phosphate buffered saline (PBS), snap-frozen in liquid nitrogen, and stored at −80°C. Using a pestle, samples were ground over liquid nitrogen to create a powder. While on ice, stages of buffer RLT (Qiagen, Valencia, CA) were added and mixed by pipetting. Following 10 min incubation, samples were centrifuged. Supernatant was loaded onto AllPrep DNA spin columns and the AllPrep DNA/RNA Mini Kit Protocol (Qiagen, Valencia, CA) was followed to isolate DNA and RNA from the same tissue sample. Total RNA was suspended in 100 μL of RNase-free water. RNA quantification was determined by NanoDrop 2000c Spectrophotometer (Thermo Fisher Scientific, Wilmington, DE).

### RNA-Seq

RNA samples were sent on dry ice to The Centre for Applied Genomics (TCAG) (The Hospital for Sick Children, Toronto, Ontario, Canada). RNA quality was analyzed by Agilent Bioanalyzer 2100 RNA Nano chip (Agilent Technologies, Santa Clara, CA) by assessing 28S/18S ratios of ribosomal RNA bands. Samples used all had RNA Integrity Numbers (RINs) greater than 8, indicating they were not degraded. RNA Library preparation followed Illumina TruSeq Stranded total RNA Library Preparation protocol to include poly(A) mRNA and lncRNA using 400 ng total RNA as starting material, with rRNA depletion using RiboZero Gold. Following RNA fragmenting at 94°C for 4 min, it was converted to double stranded cDNA, end-repaired, and 3’ adenylated for ligation of TruSeq adapters. Sample fragments were amplified with different barcode adapters for multiplex sequencing under the following PCR conditions: 10 s denaturation at 98°C, 13 cycles of 10 s at 98°C, 30 s at 60°C, 30 s at 72°C, and final extension of 5 min at 72°C. To confirm fragment size, 1 μL of final RNA library was loaded on a Bioanalyzer 2100 DNA High Sensitivity chip (Agilent Technologies, Santa Clara, CA). Kapa Library Quantification Illumina/ABI Prism Kit protocol (KAPA Biosystems) was used to quantify RNA libraries by qPCR. After pooling in equimolar amounts, libraries were paired-end sequenced on an Illumina HiSeq 2500 platform using a High Throughput Run Mode flowcell and V4 sequencing chemistry following recommended Illumina protocol to generate 126-bp long paired-end reads.

### Pseudoalignment and differential expression analysis

Paired-end reads for each sample were quantified via pseudoalignment to version 38 of the Ensembl annotation of the mouse transcriptome using kallisto (Bray et al., 2016). To estimate inferential variance of transcript abundance, 100 bootstrap samples were taken. Differential analysis of gene expression was determined using sleuth (Pimentel et al., 2017) via transcript *p*-value aggregation (Yi et al., 2018). The sleuth object model was defined based on treatment group, with Wald Tests performed to compare each experimental treatment group (ethanol, stress, and ethanol + stress) to the control group. Generalized hypergeometric tests for enrichment of GO terms and KEGG pathways were used for genes represented by transcripts differentially expressed in each treatment group using the goana and kegga functions in the *limma* software package (Ritchie et al., 2015) in R, filtered by significance (p < 0.05).

### Weighted gene co-expression network analysis

Transcript abundance in transcripts per million (tpm) was used for weighted gene co-expression network analysis (WGCNA) for transcripts detected in all samples (92 187) (Langfelder and Horvath, 2008). As less than 20 samples were used, the soft power threshold was set at 9 for production of adjacency matrices from correlation values for each combination of transcripts. A topological overlap matrix and topological overlap dissimilarity matrix were produced and used for agglomerative hierarchical clustering by average linkage method. Transcripts were clustered based on the topological overlap between them. Modules were defined using a blockwise network analysis with a maximum block size of 15 000, minimum module size of 30, merge cut height of 0.35, with deepSplit at the default 2 for medium sensitivity. Transcripts that did not cluster in a specific module were placed in module 0 and not considered for further analysis. Modules were numerically labeled by module size, with module 1 being the largest module. Each module was represented by the module eigengene (ME), the first principle component of the module.

Co-expression modules were related to 11 traits based on treatment as well as phenotypic results as previously described (Alberry and Singh, 2016). Briefly, this includes prenatal ethanol treatment, postnatal stress treatment, experimental group, Barnes maze learning score, weight at postnatal day 21, activity, distance travelled, latency to enter center zone, and number of entries into center zone of the open field test, as well as activity, and number of rears in the home cage activity test. Each of these traits was experimentally assigned as a treatment or a measured outcome significantly different following at least one of the treatments. Module-trait correlations were filtered by significance (p < 0.05). Modules of interest were selected as significantly correlated modules to each trait, resulting in 20 modules of interest. Generalized hypergeometric tests for enrichment of GO terms were used for genes represented by transcripts present in each module using Enrichr (Kuleshov et al., 2016), filtered by significance (p < 0.05). Similarly, KEGG pathways were dertermined using the kegga function in the *limma* software package (Ritchie et al., 2015) in R, filtered by significance (p < 0.05).

### qPCR for gene expression

Purified hippocampal RNA was converted to cDNA using the SuperScript IV VILO Master Mix with ezDNase Enzyme following manufacturer’s instructions (Thermo Fisher Scientific). TaqMan Assays were used to investigate the gene of interest, *Polr2a* (ID Mm00839502_m1, FAM labelled) in a multiplex reaction with TATA box binding protein (*Tbp*) as an endogenous reference gene (ID Mm01277042_m1, VIC labelled) with the TaqMan Fast Advanced Master Mix according to the manufacturer’s instructions (Applied Biosystems). The 2^−ΔΔCt^ method was used to assess relative quantity.

## ACKNOWLEDGEMENTS

The authors would like to acknowledge Eric Chater-Diehl and Benjamin Laufer for assistance with experimental design, training, and manuscript guidance, as well as undergraduate researchers Shruthi Rethi, Ali Pensamiento, David Seok, and Yuchen Li for data collection.

## DATA AVAILABILITY

Sequence data is available at GEO, accession number pending.

## SUPPLEMENTARY MATERIAL

**Supplementary Figure 1. Module creation by weighted gene co-expression network analysis.** (A) Sample clustering to detect outliers. (B) Connectivity analysis of the scale free topology fit for different soft-thresholding powers where numbers indicate the soft-thresholding power (C) mean connectivity of the network for different soft-thresholding powers, a soft-threshold of 9 was used here. (D, E, F, G, H, I, J, K) Transcript similarity clustering dendrograms for blockwise analysis for blocks 1-8, respectively, by clustering of transcripts based on topological overlap with different modules indicated by color below.

**Supplementary Table 1.** Correlation coefficients and p-values of module-trait associations for each of the 44 modules produced by WGCNA and 11 traits.

**Supplementary Table 2.** Genes implicated by transcripts in Module 19 associated with experimental treatment group and number of center zone entries in the open field test; GO terms and KEGG pathways overrepresented (p < 0.05) in module 19.

**Supplementary Table 3.** Differentially expressed gene lists for Ethanol, Stress, and Ethanol + Stress as compared to controls filtered by significance (p < 0.05); GO terms and KEGG pathways overrepresented (p < 0.05) by transcripts significantly differentially expressed (p < 0.01) for each comparison.

## REFERENCES

Ables, J. L., Breunig, J. J., Eisch, A. J. and Rakic, P. (2011). Not(ch) just development: Notch signalling in the adult brain. Nat. Rev. Neurosci. 12, 269–283.

Alberry, B. and Singh, S. M. (2016). Developmental and behavioral consequences of early life maternal separation stress in a mouse model of fetal alcohol spectrum disorder. Behav. Brain Res. 308, 94–103.

Allan, A. M., Chynoweth, J., Tyler, L. A. and Caldwell, K. K. (2003). A mouse model of prenatal ethanol exposure using a voluntary drinking paradigm. Alcohol. Clin. Exp. Res. 27, 2009–2016.

Ayata, P., Badimon, A., Strasburger, H. J., Duff, M. K., Montgomery, S. E., Loh, Y. H. E., Ebert, A., Pimenova, A. A., Ramirez, B. R., Chan, A. T., et al. (2018). Epigenetic regulation of brain region-specific microglia clearance activity. Nat. Neurosci. 21, 1049–1060.

Beech, R. D., Leffert, J. J., Lin, A., Hong, K. A., Hansen, J., Umlauf, S., Mane, S., Zhao, H. and Sinha, R. (2014). Stress-related alcohol consumption in heavy drinkers correlates with expression of miR-10a, miR-21, and components of the TAR-RNA-binding protein-associated complex. Alcohol. Clin. Exp. Res. 38, 2743–2753.

Benner, S., Endo, T., Endo, N., Kakeyama, M. and Tohyama, C. (2014). Early deprivation induces competitive subordinance in C57BL/6 male mice. Physiol. Behav. 137, 42–52.

Bray, N. L., Pimentel, H., Melsted, P. and Pachter, L. (2016). Near-optimal probabilistic RNA-seq quantification. Nat. Biotechnol. 34, 525–7.

Brose, K., Bland, K. S., Kuan, H. W., Arnott, D., Henzel, W., Goodman, C. S., Tessier-Lavigne, M. and Kidd, T. (1999). Slit proteins bind robo receptors and have an evolutionarily conserved role in repulsive axon guidance. Cell 96, 795–806.

Chastain, L. G., Franklin, T., Gangisetty, O., Cabrera, M. A., Mukherjee, S., Shrivastava, P., Jabbar, S. and Sarkar, D. K. (2019). Early life alcohol exposure primes hypothalamic microglia to later-life hypersensitivity to immune stress: possible epigenetic mechanism. Neuropsychopharmacology 1.

Chater-Diehl, E. J., Laufer, B. I., Castellani, C. A., Alberry, B. L. and Singh, S. M. (2016). Alteration of gene expression, DNA methylation, and histone methylation in free radical scavenging networks in adult mouse hippocampus following fetal alcohol exposure. PLoS One 11, e0154836.

Chater-Diehl, E. J., Laufer, B. I. and Singh, S. M. (2017). Changes to histone modifications following prenatal alcohol exposure: An emerging picture. Alcohol 60, 41–52.

Chokroborty-Hoque, A., Alberry, B. and Singh, S. M. (2014). Exploring the complexity of intellectual disability in fetal alcohol spectrum disorders. Front. Pediatr. 2,.

Choleris, E., Thomas, A. W., Kavaliers, M. and Prato, F. S. (2001). A detailed ethological analysis of the mouse open field test: Effects of diazepam, chlordiazepoxide and an extremely low frequency pulsed magnetic field. Neurosci. Biobehav. Rev. 25, 235–260.

Chudley, A. E., Conry, J., Cook, J. L., Loock, C., Rosales, T., LeBlanc, N. and Disorder, P. H. A. of C. N. A. C. on F. A. S. (2005). Fetal alcohol spectrum disorder: Canadian guidelines for diagnosis. Can. Med. Assoc. J. 172, S1–S21.

Coggins, T. E., Timler, G. R. and Olswang, L. B. (2007). A State of Double Jeopardy: Impact of Prenatal Alcohol Exposure and Adverse Environments on the Social Communicative Abilities of School-Age Children With Fetal Alcohol Spectrum Disorder. Lang. Speech. Hear. Serv. Sch. 38, 117–127.

Cornman-Homonoff, J., Kuehn, D., Aros, S., Carter, T. C., Conley, M. R., Troendle, J., Cassorla, F. and Mills, J. L. (2012). Heavy prenatal alcohol exposure and risk of stillbirth and preterm delivery. J. Matern. Neonatal Med. 25, 860–863.

Cuthbert, P. C., Stanford, L. E., Coba, M. P., Ainge, J. A., Fink, A. E., Opazo, P., Delgado, J. Y., Komiyama, N. H., O’Dell, T. J. and Grant, S. G. N. (2007). Synapse-Associated Protein 102/dlgh3 Couples the NMDA Receptor to Specific Plasticity Pathways and Learning Strategies. J. Neurosci. 27, 2673–2682.

Duric, V., Banasr, M., Licznerski, P., Schmidt, H. D., Stockmeier, C. A., Simen, A. A., Newton, S. S. and Duman, R. S. (2010). A negative regulator of MAP kinase causes depressive behavior. Nat. Med. 16, 1328–1332.

Fenoglio, K. A., Brunson, K. L. and Baram, T. Z. (2006). Hippocampal neuroplasticity induced by early-life stress: functional and molecular aspects. Front. Neuroendocrinol. 27, 180–192.

Franklin, T. B., Russig, H., Weiss, I. C., Graff, J., Linder, N., Michalon, A., Vizi, S. and Mansuy, I. M. (2010). Epigenetic transmission of the impact of early stress across generations. Biol. Psychiatry 68, 408–415.

Fromer, M., Roussos, P., Sieberts, S. K., Johnson, J. S., Kavanagh, D. H., Perumal, T. M., Ruderfer, D. M., Oh, E. C., Topol, A., Shah, H. R., et al. (2016). Gene expression elucidates functional impact of polygenic risk for schizophrenia. Nat. Neurosci. 19, 1442–1453.

Hatalski, C. G., Brunson, K. L., Tantayanubutr, B., Chen, Y. and Baram, T. Z. (2000). Neuronal activity and stress differentially regulate hippocampal and hypothalamic corticotropin-releasing hormone expression in the immature rat. Neuroscience 101, 571–580.

Henry, J., Sloane, M. and Black-Pond, C. (2007). Neurobiology and Neurodevelopmental Impact of Childhood Traumatic Stress and Prenatal Alcohol Exposure. Lang. Speech. Hear. Serv. Sch. 38, 99–108.

Hill, A. S., Sahay, A. and Hen, R. (2015). Increasing Adult Hippocampal Neurogenesis is Sufficient to Reduce Anxiety and Depression-Like Behaviors. Neuropsychopharmacology 40, 2368–2378.

Juul, S. E., Beyer, R. P., Bammler, T. K., Farin, F. M. and Gleason, C. A. (2011). Effects of neonatal stress and morphine on murine hippocampal gene expression. Pediatr. Res. 69, 285–292.

Kaminen-Ahola, N., Ahola, A., Flatscher-Bader, T., Wilkins, S. J., Anderson, G. J., Whitelaw, E. and Chong, S. (2010). Postnatal growth restriction and gene expression changes in a mouse model of fetal alcohol syndrome. Birth defects Res. A, Clin. Mol. Teratol. 88, 818–826.

Kesmodel, U., Wisborg, K., Olsen, S. F., Henriksen, T. B. and Secher, N. J. (2002). Moderate alcohol intake in pregnancy and the risk of spontaneous abortion. Alcohol Alcohol 37, 87–92.

Khalid, O., Kim, J. J., Kim, H.-S. S., Hoang, M., Tu, T. G., Elie, O., Lee, C., Vu, C., Horvath, S., Spigelman, I., et al. (2014). Gene expression signatures affected by alcohol-induced DNA methylomic deregulation in human embryonic stem cells. Stem Cell Res. 12, 791–806.

Kisely, S., Abajobir, A. A., Mills, R., Strathearn, L., Clavarino, A. and Najman, J. M. (2018). Child maltreatment and mental health problems in adulthood: Birth cohort study. Br. J. Psychiatry 213, 698–703.

Kleiber, M. L., Wright, E. and Singh, S. M. (2011). Maternal voluntary drinking in C57BL/6J mice: advancing a model for fetal alcohol spectrum disorders. Behav. Brain Res. 223, 376–387.

Koponen, A. M., Kalland, M. and Autti-Rämö, I. (2009). Caregiving environment and socioemotional development of foster-placed FASD-children. Child. Youth Serv. Rev. 31, 1049–1056.

Koponen, A. M., Kalland, M., Autti-Rämö, I., Laamanen, R. and Suominen, S. (2013). Socioemotional development of children with foetal alcohol spectrum disorders in long-term foster family care: a qualitative study. Nord. Soc. Work Res. 3, 38–58.

Kreisel, T., Frank, M. G., Licht, T., Reshef, R., Ben-Menachem-Zidon, O., Baratta, M. V, Maier, S. F. and Yirmiya, R. (2014). Dynamic microglial alterations underlie stress-induced depressive-like behavior and suppressed neurogenesis. Mol. Psychiatry 19, 699–709.

Kuleshov, M. V, Jones, M. R., Rouillard, A. D., Fernandez, N. F., Duan, Q., Wang, Z., Koplev, S., Jenkins, S. L., Jagodnik, K. M., Lachmann, A., et al. (2016). Enrichr: a comprehensive gene set enrichment analysis web server 2016 update. Nucleic Acids Res. 44, W90–7.

Lange, S., Shield, K., Rehm, J. and Popova, S. (2013). Prevalence of fetal alcohol spectrum disorders in child care settings: a meta-analysis. Pediatrics 132, e980–95.

Lange, S., Probst, C., Gmel, G., Rehm, J., Burd, L. and Popova, S. (2017). Global prevalence of fetal alcohol spectrum disorder among children and youth: A systematic review and meta-analysis. JAMA Pediatr. 171, 948–956.

Langfelder, P. and Horvath, S. (2008). WGCNA: an R package for weighted correlation network analysis. BMC Bioinformatics 9, 559.

Liu, Z., Niu, Y., Xie, M., Bu, Y., Yao, Z. and Gao, C. (2014). Gene expression profiling analysis reveals that DLG3 is down-regulated in glioblastoma. J. Neurooncol. 116, 465–476.

Lombard, Z., Tiffin, N., Hofmann, O., Bajic, V. B., Hide, W. and Ramsay, M. (2007). Computational selection and prioritization of candidate genes for Fetal Alcohol Syndrome. BMC Genomics 8, 389.

Louis, L. K., Gopurappilly, R., Surendran, H., Dutta, S. and Pal, R. (2018). Transcriptional profiling of human neural precursors post alcohol exposure reveals impaired neurogenesis via dysregulation of ERK signaling and miR-145. J. Neurochem. 146, 47–62.

Marjonen, H., Sierra, A., Nyman, A., Rogojin, V., Gröhn, O., Linden, A.-M. M., Hautaniemi, S., Kaminen-Ahola, N., Grohn, O., Linden, A.-M. M., et al. (2015). Early maternal alcohol consumption alters hippocampal DNA methylation, gene expression and volume in a mouse model. PLoS One 10, e0124931.

May, P. A., Chambers, C. D., Kalberg, W. O., Zellner, J., Feldman, H., Buckley, D., Kopald, D., Hasken, J. M., Xu, R., Honerkamp-Smith, G., et al. (2018). Prevalence of fetal alcohol spectrum disorders in 4 US communities. JAMA - J. Am. Med. Assoc. 319, 474–482.

Muralidharan, P., Sarmah, S. and Marrs, J. A. (2018). Retinal Wnt signaling defect in a zebrafish fetal alcohol spectrum disorder model. PLoS One 13, e0201659.

Ninh, V. K., El Hajj, E. C., Mouton, A. J. and Gardner, J. D. (2019). Prenatal Alcohol Exposure Causes Adverse Cardiac Extracellular Matrix Changes and Dysfunction in Neonatal Mice. Cardiovasc. Toxicol. 1–12.

Oomen, C. A., Soeters, H., Audureau, N., Vermunt, L., van Hasselt, F. N., Manders, E. M. M., Joëls, M., Lucassen, P. J. and Krugers, H. (2010). Severe early life stress hampers spatial learning and neurogenesis, but improves hippocampal synaptic plasticity and emotional learning under high-stress conditions in adulthood. J. Neurosci. 30, 6635–45.

Peters, A. H. F. M., O’Carroll, D., Scherthan, H., Mechtler, K., Sauer, S., Schöfer, C., Weipoltshammer, K., Pagani, M., Lachner, M., Kohlmaier, A., et al. (2001). Loss of the Suv39h histone methyltransferases impairs mammalian heterochromatin and genome stability. Cell 107, 323–337.

Pillai, A. G., Arp, M., Velzing, E., Lesuis, S. L., Schmidt, M. V., Holsboer, F., Joëls, M. and Krugers, H. J. (2018). Early life stress determines the effects of glucocorticoids and stress on hippocampal function: Electrophysiological and behavioral evidence respectively. Neuropharmacology 133, 307–318.

Pimentel, H., Bray, N. L., Puente, S., Melsted, P. and Pachter, L. (2017). Differential analysis of RNA-seq incorporating quantification uncertainty. Nat. Methods 14, 687–690.

Popova, S., Lange, S., Burd, L. and Rehm, J. (2016). The economic burden of fetal alcohol spectrum disorder in Canada in 2013. Alcohol Alcohol. 51, 367–375.

Popova, S., Lange, S., Probst, C., Gmel, G. and Rehm, J. (2017). Global Prevalence of Alcohol Use and Binge Drinking During Pregnancy and Fetal Alcohol Spectrum Disorder. Biochem. Cell Biol. bcb-2017-0077.

Popova, S., Lange, S., Chudley, A. E., Reynolds, J. N., Rehm, J., May, P. A. and Riley, E. P. (2018). World Health Organization International Study on the Prevalence of Fetal Alcohol Spectrum Disorder (FASD). Cent. Addit. Ment. Heal.

Price, A., Cook, P. A., Norgate, S. and Mukherjee, R. (2017). Prenatal alcohol exposure and traumatic childhood experiences: A systematic review. Neurosci. Biobehav. Rev. 80, 89–98.

Prut, L. and Belzung, C. (2003). The open field as a paradigm to measure the effects of drugs on anxiety-like behaviors: A review. Eur. J. Pharmacol. 463, 3–33.

Radulescu, E., Jaffe, A. E., Straub, R. E., Chen, Q., Shin, J. H., Hyde, T. M., Kleinman, J. E. and Weinberger, D. R. (2018). Identification and prioritization of gene sets associated with schizophrenia risk by co-expression network analysis in human brain. Mol. Psychiatry 1.

Rice, C. J., Sandman, C. A., Lenjavi, M. R. and Baram, T. Z. (2008). A novel mouse model for acute and long-lasting consequences of early life stress. Endocrinology 149, 4892–4900.

Ritchie, M. E., Phipson, B., Wu, D., Hu, Y., Law, C. W., Shi, W. and Smyth, G. K. (2015). limma powers differential expression analyses for RNA-sequencing and microarray studies. Nucleic Acids Res. 43, e47.

Romeo, R. D., Mueller, A., Sisti, H. M., Ogawa, S., McEwen, B. S. and Brake, W. G. (2003). Anxiety and fear behaviors in adult male and female C57BL/6 mice are modulated by maternal separation. Horm. Behav. 43, 561–567.

Rosenberg, M. J., Wolff, C. R., El-Emawy, A., Staples, M. C., Perrone-Bizzozero, N. I. and Savage, D. D. (2010). Effects of moderate drinking during pregnancy on placental gene expression. Alcohol 44, 673–690.

Sabra, S., Malmqvist, E., Almeida, L., Gratacos, E. and Gomez Roig, M. D. (2018). Differential correlations between maternal hair levels of tobacco and alcohol with fetal growth restriction clinical subtypes. Alcohol 70, 43–49.

Savignac, H. M., Dinan, T. G. and Cryan, J. F. (2011). Resistance to early-life stress in mice: effects of genetic background and stress duration. Front. Behav. Neurosci. 5, 13.

Schneider, J. S., Anderson, D. W., Talsania, K., Mettil, W. and Vadigepalli, R. (2012). Effects of developmental lead exposure on the hippocampal transcriptome: Influences of sex, developmental period, and lead exposure level. Toxicol. Sci. 129, 108–125.

Sokol, R. J., Janisse, J. J., Louis, J. M., Bailey, B. N., Ager, J., Jacobson, S. W. and Jacobson, J. L. (2007). Extreme prematurity: An alcohol-related birth effect. Alcohol. Clin. Exp. Res. 31, 1031–1037.

Spijker, S. (2011). Dissection of Rodent Brain Regions. In Neuroproteomics (ed. Li, W. W. K.), pp. 13–26. Humana Press.

Stankiewicz, A. M., Goscik, J., Swiergiel, A. H., Majewska, A., Wieczorek, M., Juszczak, G. R., Lisowski, P. A.M. S. J. G. A.H. S., et al. (2014). Social stress increases expression of hemoglobin genes in mouse prefrontal cortex. BMC Neurosci. 15, 130.

Tarpey, P., Parnau, J., Blow, M., Woffendin, H., Bignell, G., Cox, C., Cox, J., Davies, H., Edkins, S., Holden, S., et al. (2004). Mutations in the DLG3 Gene Cause Nonsyndromic X-Linked Mental Retardation. Am. J. Hum. Genet. 75, 318–324.

Tynan, R. J., Naicker, S., Hinwood, M., Nalivaiko, E., Buller, K. M., Pow, D. V., Day, T. A. and Walker, F. R. (2010). Chronic stress alters the density and morphology of microglia in a subset of stress-responsive brain regions. Brain. Behav. Immun. 24, 1058–1068.

Urrutia, R. (2003). KRAB-containing zinc-finger repressor proteins. Genome Biol. 4, 231.

Veenema, A. H., Reber, S. O., Selch, S., Obermeier, F. and Neumann, I. D. (2008). Early life stress enhances the vulnerability to chronic psychosocial stress and experimental colitis in adult mice. Endocrinology 149, 2727–36.

Watanabe, Y., Miyasaka, K. Y., Kubo, A., Kida, Y. S., Nakagawa, O., Hirate, Y., Sasaki, H. and Ogura, T. (2017). Notch and Hippo signaling converge on Strawberry Notch 1 (Sbno1) to synergistically activate Cdx2 during specification of the trophectoderm. Sci. Rep. 7, 46135.

Wei, Q., Fentress, H. M., Hoversten, M. T., Zhang, L., Hebda-Bauer, E. K., Watson, S. J., Seasholtz, A. F. and Akil, H. (2012). Early-life forebrain glucocorticoid receptor overexpression increases anxiety behavior and cocaine sensitization. Biol. Psychiatry 71, 224–231.

Winckelmans, E., Vrijens, K., Tsamou, M., Janssen, B. G., Saenen, N. D., Roels, H. A., Kleinjans, J., Lefebvre, W., Vanpoucke, C., De Kok, T. M., et al. (2017). Newborn sex-specific transcriptome signatures and gestational exposure to fine particles: findings from the ENVIRONAGE birth cohort. Environ. Heal. A Glob. Access Sci. Source 16, 52.

Yi, L., Pimentel, H., Bray, N. L. and Pachter, L. (2018). Gene-level differential analysis at transcript-level resolution. Genome Biol. 19, 53.

Yuan, F., Chen, X., Liu, J., Feng, W., Wu, X. and Chen, S. yu (2017). Up-regulation of Siah1 by ethanol triggers apoptosis in neural crest cells through p38 MAPK-mediated activation of p53 signaling pathway. Arch. Toxicol. 91, 775–784.

