## Supplementary figures and images for "Hippocampal transcriptome analysis following maternal separation implicates altered RNA processing in a mouse model of fetal alcohol spectrum disorder"

### Supplementary Figure 1

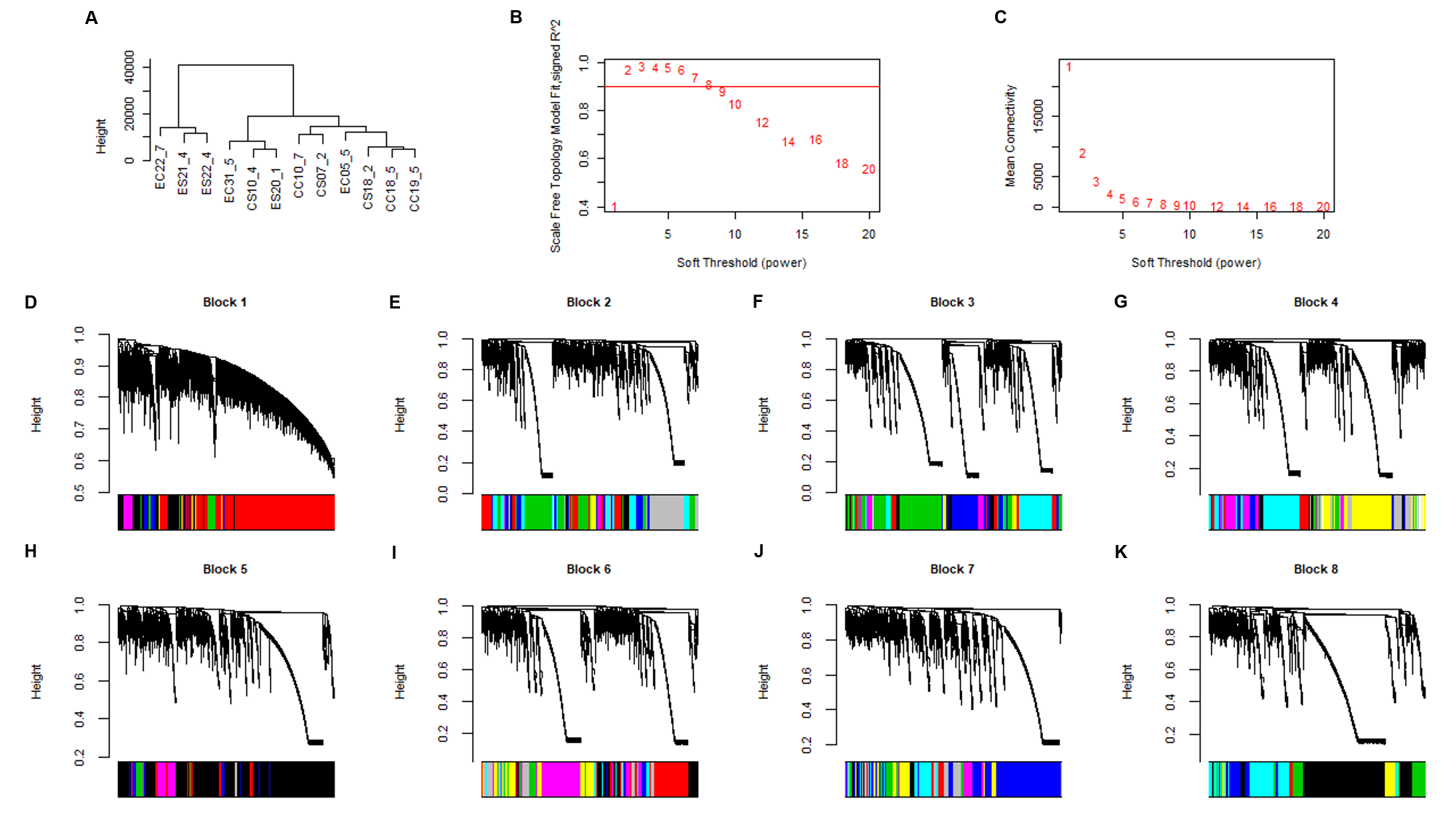
